# Expectations boost the reconstruction of auditory features from electrophysiological responses to noisy speech

**DOI:** 10.1101/2021.09.06.459160

**Authors:** Andrew W. Corcoran, Ricardo Perera, Matthieu Koroma, Sid Kouider, Jakob Hohwy, Thomas Andrillon

## Abstract

Online speech processing imposes significant computational demands on the listening brain, the underlying mechanisms of which remain poorly understood. Here, we exploit the perceptual ‘pop-out’ phenomenon (i.e. the dramatic improvement of speech intelligibility after receiving information about speech content) to investigate the neurophysiological effects of prior expectations on degraded speech comprehension. We recorded electroencephalography and pupillometry from 21 adults while they rated the clarity of noise-vocoded and sine-wave synthesised sentences. Pop-out was reliably elicited following visual presentation of the corresponding written sentence, but not following incongruent or neutral text. Pop-out was associated with improved reconstruction of the acoustic stimulus envelope from low-frequency EEG activity, implying that improvements in perceptual clarity were mediated via top-down signals that enhance the quality of cortical speech representations. Spectral analysis further revealed that pop-out was accompanied by a reduction in theta-band power, consistent with predictive coding accounts of acoustic filling-in and incremental sentence processing. Moreover, delta-band power, alpha-band power, and pupil diameter were all increased following the provision of *any* written sentence information, irrespective of content. Together, these findings reveal distinctive profiles of neurophysiological activity that differentiate the content-specific processes associated with degraded speech comprehension from the context-specific processes invoked under adverse listening conditions.

## Introduction

The ability to understand spoken language is a remarkable feat of human cognition. Fluent speech recognition requires the parsing of a continuously changing acoustic signal into a series of discrete units, and the mapping of these units onto abstract representations spanning multiple scales (Halle and Stevens 1962; Hickok and Poeppel 2007). Such processing must occur quickly enough to keep abreast of the unfolding speech stream (Christiansen and Chater 2016), while remaining robust to signal variation and degradation (Mattys et al. 2012; Guediche et al. 2014). There is growing consensus that the brain meets these demands by predicting sensory input on the basis of prior knowledge (Kuperberg and Jaeger 2016; Bornkessel-Schlesewsky and Schlesewsky 2019; Brodbeck and Simon 2020). However, the neurocomputational mechanisms supporting such processes remain poorly understood.

Prediction has long been accorded an important role in language comprehension (e.g., Miller and Isard, 1963; Tulving and Gold, 1963). Contemporary *predictive coding* models of speech processing formalise this notion in terms of (hierarchical) Bayesian inference, whereby perceptual experience reflects the integration of ‘top-down’ prior expectations (derived, e.g., from lexical, speaker, or world-knowledge) and ‘bottom-up’ sensory evidence (see Friston and Kiebel, 2009; Heilbron and Chait, 2018). On this view, resolved speech content constitutes the brain’s ‘best guess’ about the causes of its sensory input, given an internal model of the way sensations are generated (cf. ‘analysis-by-synthesis’; Halle and Stevens, 1959; Poeppel et al., 2008).

Prior knowledge plays a decisive role in word recognition. Under adverse listening conditions, speech intelligibility can be improved by the provision of prior information about lexical content (e.g., hearing the undistorted version of the word or reading its written transcription; Giraud et al., 2004; Dehaene-Lambertz et al., 2005). Such information typically engenders a dramatic improvement in the subjective clarity of the degraded utterance -- a striking change in perceptual experience referred to as ‘pop-out’ (Davis et al. 2005).

### Predictive coding and perceptual pop-out

Consistent with the empirical predictions of predictive coding theory, auditory cortical responses to degraded words tend to be suppressed during the experience of pop-out (Sohoglu et al., 2012; Sohoglu and Davis, 2016; cf. Banellis et al., 2020). Similar findings have been observed during the restoration or ‘filling-in’ of speech sounds at the sublexical level (Riecke et al. 2012; Shahin et al. 2012; Leonard et al. 2016). Interestingly, auditory cortical responses to degraded words depend on both the severity of stimulus degradation and the accuracy of prior expectations: while clearer speech evokes greater suppression of cortical responses when expectations are realised, neural activity is *enhanced* when expectations are violated (Blank and Davis 2016; Sohoglu and Davis 2020). This coheres with the view that the discrepancy between predicted and actual sensory inputs (*prediction error*) is modulated by the quality of sensory evidence, whereby noisier stimuli are assigned a lower degree of confidence or *precision*.

Few studies have investigated the predictive mechanisms of pop-out during continuous speech processing. Consistent with the (sub)lexical studies mentioned above, functional magnetic resonance imaging has shown that degraded sentences elicit increased primary auditory cortical activation relative to clear speech, indicative of increased sensory prediction error (Tuennerhoff and Noppeney 2016). The contrast between unintelligible and intelligible speech was characterised by the activation of higher-order cortical regions that appeared to modulate lower-level sensory processing. Electrocorticographic recordings of high-frequency broadband activity have further revealed the rapid tuning of auditory cortical ensembles during sentence pop-out, whereby hearing the undistorted sentence renders neuronal populations more sensitive to speech-specific spectro-temporal auditory features of the subsequently-replayed degraded stimulus (Holdgraf et al., 2016). These findings suggest prior exposure to clear speech induces altered patterns of neural activity that serve to enhance the extraction of linguistic information from noisy input. However, this study was unable to establish whether such cortical plasticity is driven by top-down or bottom-up mechanisms.

More recently, ‘cortical tracking’ techniques (Wöstmann, Fiedler, et al. 2017; Beier et al. 2021) have been used to investigate electroencephalographic (EEG) correlates of degraded speech comprehension. Baltzell and colleagues (2017) reported that the cross-correlation between the amplitude envelopes of the acoustic stimulus and broadband (1-50 Hz) EEG activity is increased while listening to degraded vs. clear sentences. This effect was amplified when degraded stimuli were rendered intelligible by prior exposure to the undistorted version of the utterance, implying that prior experience of sentence content improves the alignment (or ‘entrainment’; see Obleser and Kayser 2019) of neural oscillations to the speech envelope. Di Liberto and colleagues (2018) further demonstrated that sentence pop-out is associated with enhanced phonemic encoding activity in the delta (but not the theta) band. This effect was accompanied by evidence of an overall reduction in phoneme-level processing, concordant with the predictive coding view that accurate prior expectations suppress low-level cortical responses to sensory input.

### The present study

In this study, we deploy a unique combination of analytic techniques spanning stimulus reconstruction, time-frequency/spectral analysis, and pupillometry to bring recent findings from the speech pop-out literature into contact with the spectral architecture of language processing and perceptual inference. Specifically, this multimodal suite of analyses was designed to complement previous model-based analyses of degraded speech processing (e.g., Holdgraf et al., 2016; Di Liberto et al., 2018; Sohoglu and Davis, 2020) with mechanistic insights into the neurophysiological activity that accompanies pop-out. While these studies showed that prior information led to enhanced sensory processing and more precise neural tuning to acoustic features, our use of backward modelling (i.e. stimulus reconstruction) allowed us to directly quantify changes in the fidelity of auditory cortical speech representations depending on the availability of prior information (Cervantes Constantino and Simon 2018), and to map these effects to their neurophysiological correlates.

We furthermore compared the effects of prior information on sentence processing across two complementary forms of speech degradation. To do so, we used: (1) *noise-vocoding*, which obscures spectral cues with white noise (Shannon et al., 1995); and (2) *sine-wave synthesis*, which obliterates fine-grained temporal structure (Remez et al., 1981). Although vocoding is a popular technique for degrading speech stimuli, little is known about the neurophysiological correlates of pop-out in sine-wave speech (but see Lee and Noppeney 2011; Khoshkhoo et al. 2018). Crucially, we trained our decoder on an independent dataset (EEG recorded while listening to an undistorted narrative) to test our prediction that stimulus reconstruction tracks the enhanced neural representation of sensory content as opposed to specific features of degraded speech.

Finally, our study departs from previous reports by rigorously controlling the extent to which differences in sensory processing and neurophysiological activity can be ascribed to top-down mechanisms. Given the exquisite sensitivity of auditory cortical ensembles to spectro-temporal speech features (cf. Holdgraf et al., 2016), exposure to clear speech might induce bottom-up changes in cortical activity that facilitate subsequent encoding of the degraded stimulus. The present study avoids this potential confound by using visual information (written text) to instil top-down expectations about the linguistic content of degraded stimuli (Wild et al. 2012; Sohoglu et al. 2014). We also contrasted the effects of correct and incorrect prior information against a baseline condition in which no written sentence information was provided, thus enabling us to characterise the influence of prior information on auditory and higher-level processing independent of its lexical content, while also disentangling the effects of ‘misinformed’ vs. ‘uninformed’ expectations. In this way, we were able to cross-modally manipulate prior knowledge and sentence intelligibility while holding prior exposure to acoustic stimuli constant across conditions.

We hypothesised that a multivariate decoder model trained on EEG responses to undistorted continuous speech would reconstruct the acoustic envelope of degraded sentences more accurately after listeners had been provided with correct (but not incorrect or no) written information about sentence content. This prediction was borne out in both the noise-vocoded and sine-wave speech conditions. Moreover, this effect was accompanied by a selective reduction in theta-band activity. Our analysis also revealed general effects of prior expectation, whereby delta-band power, alpha-band power, and pupil diameter were all increased while the listener evaluated whether degraded speech corresponded to written sentence information. Together, these results demonstrate that top-down mechanisms shape the online integration of bottom-up sensory information during continuous speech processing.

## Materials and methods

### Participants

Twenty-one native English-speaking adults were recruited to participate in this study. Of these, two were excluded due to faulty EEG recordings. The remaining sample comprised 8 females and 11 males aged 19 to 33 years (*M* = 25.8, *SD* = 4.5). All participants reported normal (or corrected-to-normal) vision and audition.

All participants provided written, informed consent, and were remunerated AU$30 for their time. This protocol was approved by the Monash University Human Research Ethics Committee (Project ID: 10994).

### Stimuli

A total of 80 pairs of English sentences were constructed. These pairs had similar but not identical grammatical structures and lengths (11.7 words on average, range: 8-15). They were divided into 5 lists of 16 pairs (32 sentences per list). Each sentence was vocoded using Apple OS’s noise-to-speech command ‘say’ (voice: ‘Alex’; gender: male; sampling rate: 44.1 kHz; rate: 200 words/minute). Each vocoded sentence was approximately 3.5 s long and was then concatenated three times to obtain audio files of ∼10.5 s. The sounds were saved in the Audio Interchange File Format (AIFF) format and converted to the MPEG-1 Audio Layer III (MP3) format using the “Swiss Army Knife of Sound” (SoX) command line utility.

We then used publicly available scripts written for PRAAT (Boersma and Weenink 2011) to turn clear speech into sine-wave speech (SWS) and noise-vocoded speech (NVS). These files were saved in the Waveform Audio File Format (WAV) format. Clear audio files were also converted to the WAV format. In SWS, phonemes’ formants are replaced by sinusoids at the same frequency, stripping fine-grained temporal acoustic features from the original clear speech and thereby making SWS speech-like but unintelligible (Remez et al. 1981). In NVS, the envelope of clear speech in a set of fixed logarithmically-spaced frequency bands (here, 7 bands) was used to modulate the amplitude of band-limited white noise for each frequency band. This transformation preserves the temporal cues of the original signal but erases the spectral cues (Shannon et al. 1995). Consequently, SWS and NVS represent two complementary ways of degrading clear speech by removing fine-grained temporal cues (SWS) or spectral information (NVS; see Figure 1A).

**Figure 1:**
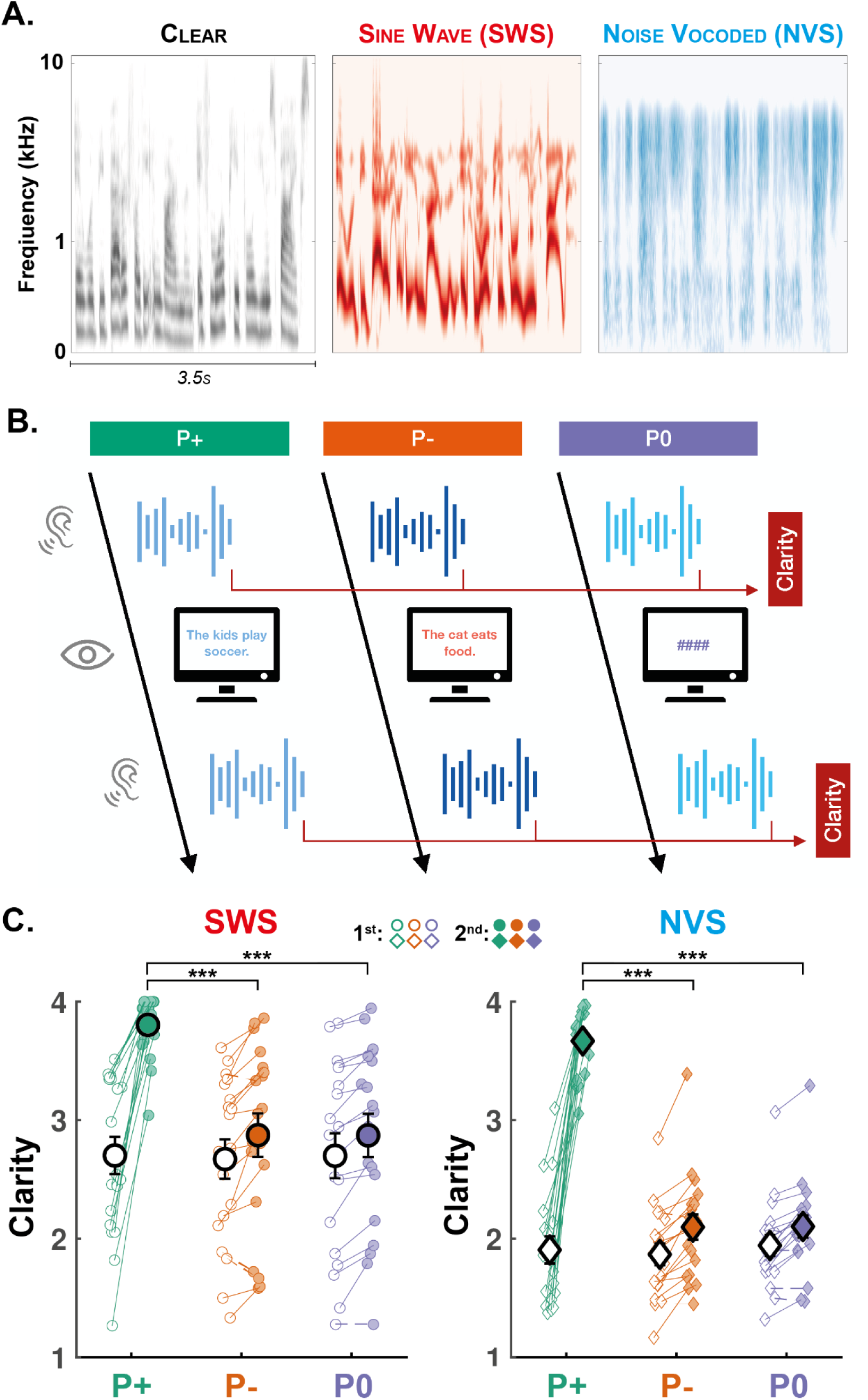
Experimental design and behavioural results. A: Cochlear representations (see Methods for details) of 3.5 s of clear speech (left), Sine Wave Speech (SWS; middle) and Noise Vocoded Speech (NVS; right). B: In each trial, participants listened to two repetitions of the same noisy speech. The two presentations of the stimuli were interleaved with either (i) the corresponding written sentence (correct prior, condition P+), (ii) a different sentence (incorrect prior, condition P-), or (iii) hash symbols (no prior, condition P0). Following each presentation of the stimulus, participants were asked to indicate the subjective clarity of the stimulus they heard. EEG was recorded throughout the task. C: Clarity ratings for the SWS (left, circles) and NVS (right, diamonds) stimuli. Participants were asked to rate the stimuli after the 1st (unfilled circles and diamonds) and 2nd (filled circles and diamonds) presentations. Clarity ratings are averaged for each stimulus type and prior condition (P+: green, P-: orange, and P0: purple). Individual data-points are shown with small circles (SWS) and diamonds (NVS). The two average ratings of each participant and each category are connected with a continuous line if it increases from the 1st to 2nd presentation and a dashed line if it decreases. Large circles and diamonds show the average across the sample (N=19 participants) and error bars show the standard error of the mean (SEM) across participants. Stars indicate the significance levels of post-hoc contrasts across condition levels (marginalised over stimulus type; ***: p<.001, **: p<.01, *: p<.05).

The amplitude of the degraded speech was equalised across all sentences and the duration was adapted to a fixed 10.5 s interval using the VSOLA algorithm. In addition to these sentences, in the training session, we also played to participants an audiobook (Cat-Skin from Grimms’ Fairy Tales, LibriVox) for a duration of 11’ 38’’. The properties of the speech (female voice, rate, etc.) were not modified except for the overall volume (same volume as degraded sentences). All auditory stimuli were delivered using the Psychtoolbox extension (v3.0.14; Brainard, 1997) for Matlab (R2018b; The MathWorks, Natick, MA, USA) running on Linux. The stimuli were played using speakers placed in front of the participant.

### Experimental design and procedure

Participants performed the experimental task in a well-lit room while sitting at a desk with their head stabilised on a chinrest ∼50 cm from the monitor. Following a 9-point eye-tracker calibration, participants were instructed to actively attend to the audiobook narration (training) while maintaining fixation on a cross at the centre of the computer screen. They subsequently performed 6 blocks of 16 experimental trials each (test trials) for a total of 96 trials (16 trials per condition). Participants were instructed to maintain central fixation and refrain from excessive blinking while listening to the sentence presentations, but were permitted to blink and saccade outside these periods. Blocks were separated by self-paced breaks, with a recalibration of the eye-tracker prior to block 4. In total, the experimental procedure lasted approximately 75 min.

Each test trial started with the presentation of one noisy stimulus (NVS or SWS; 10.5 s long). Participants were then asked to rate the clarity (intelligibility) of the noisy stimulus on a 4-point scale (1 = “I did not understand anything”; 2 = “I understood some of the sentence”; 3 = “I understood most of the sentence”; 4 = “I clearly understood everything”). Following this first clarity rating, participants were visually displayed either the corresponding written sentence (P+ condition), a different sentence (P- condition), or four hash symbols in lieu of a sentence (P0 condition), for a fixed duration of 4 s. In all cases, the same noisy stimulus was presented a second time and participants were asked to rate the clarity of the stimulus using the same 4-point scale. Following this, when a sentence was visually displayed between the two presentations (P+ and P- conditions), participants were asked to indicate whether the displayed sentence corresponded to the noisy stimulus (Yes or No). A pause of 1.5 to 2 s (random jitter) was introduced before starting the next trial. See Figure 1B for a schematic illustration of the trial procedure.

Participants heard a total of 96 sentence stimuli. Sentences were sampled from a pool of 80 stimulus pairs distributed amongst 5 lists (Lists A-E, 160 stimuli). Four lists of 16 stimulus pairs were attributed to experimental conditions (stimulus type: SWS or NVS; prior condition: P+, P-). For these conditions, only one member of each stimulus pair was presented (e.g., 16 NVS P- stimuli from list A, 16 NVS P+ from list B, 16 SWS P- from list C, 16 SWS P+ from list D; total: 64) and the remaining paired stimuli were not used. The remaining list of 16 stimulus pairs was attributed to the condition P0 (e.g., 16 NVS P0 from list E and 16 SWS P0 from list E; total: 32). The attribution was randomly assigned and counterbalanced across participants following a latin-square design. For conditions P+ and P-, the participants were exposed to one sentence per pair from the corresponding lists (see section above on Stimuli). This allowed us to present to participants, in the case of the P- condition, a sentence close to (but different from) the one heard in the trial, and never heard or seen earlier or later on in the experiment. For the P0 condition, as no pairing was needed since no written sentence was shown to the participant, the remaining list of 16 pairs was split into the SWS and NVS conditions and formed 16 stimuli per condition. Overall, every stimulus was novel when presented the first time and heard exactly twice within one trial throughout the whole experiment and across all conditions.

### EEG acquisition and preprocessing

The electroencephalogram (EEG) was continuously recorded during both the training (audiobook) and test trials (noisy speech) from 64 Ag/AgCl EasyCap mounted active electrodes. The recording was acquired at a sampling rate of 500 Hz using a BrainAmp system in conjunction with BrainVision Recorder (v1.21.0402; Brain Products GmbH, Gilching, Germany). AFz served as the ground electrode and FCz as the online reference.

Offline preprocessing was performed in MATLAB R2019b (v9.7.0.1319299; The MathWorks, Natick, MA, USA) using custom-build scripts incorporating functions from the FieldTrip (v20200623; Oostenveld et al., 2011) and EEGLAB (v2019.1; Delorme and Makeig, 2004) toolboxes. For the training data, the EEG was segmented in a single epoch from 5 s before the start of the audiobook to 5 s after its end. For test trials, EEG data were segmented into 20 s epochs beginning 5 s before stimulus onset. All epochs were centred around 0 prior to high- and low-pass filtering (1 Hz and 125 Hz, respectively; two-pass 4th-order Butterworth filters). A notch (discrete Fourier transform) filter was also applied at 50 and 100 Hz to mitigate line noise.

For test trials, epoch and channel data were manually screened for excessive artefact using the ‘ft_rejectvisual’ function. A median 3 channels (range = [1, 5]) and 2 epochs (range = [0, 5]) were rejected per participant (note, an additional 5 trials were missing for one participant due to a technical error). For training data, we performed only the channel rejection procedure. Rejected channels were interpolated via the weighted neighbour approach as implemented in the ‘ft_channelrepair’ function (where channel neighbours were defined by triangulation).

Channels were re-referenced to the common average prior to independent component analysis (‘runica’ implementation in FieldTrip of the logistic infomax ICA algorithm; Bell and Sejnowski, 1995). ICA was performed on the test and training data separately to ensure systematic differences between clear and degraded speech processing did not impair or bias source separation. Components were visually inspected and those identified as ocular (median number of rejected components = 2; range = [0, 3]), cardiac (median = 0, range = [0, 2]), or non-physiological (median = 0, range = [0, 2]) in origin were subtracted prior to backprojection.

### Pupillometry acquisition and preprocessing

Eye-movements and pupil size on both eyes were recorded with a Tobii Pro TX300 system (Tobii Pro) at a sampling rate of 300 Hz. We recorded good-quality data in only 17 participants. One participant had incomplete data (43/96 trials). The eye-tracker was calibrated at the start of each recording. Blinks were detected as interruptions of the eye-tracking signal on each eye independently (maximum duration = 5 s). For each of these blinks, the pupil size was corrected by linearly interpolating the median signal preceding the blink onset ([-0.1, 0]s) and following the blink offset ([0, 0.1]s). The corrected signal was then low-pass filtered below 6 Hz (two-pass 4th order Butterworth filter) and the pupil size for each eye averaged together. The continuous averaged pupil data were then epoched according to the presentation onset ([-1, 11]s) and both the first and second presentation windows were baseline corrected using the average pupil size before the first presentation ([-1, 0]s). Event-related pupil dilation responses were computed on these epochs (see Figure 4B).

### Data analysis

#### Stimulus reconstruction

We used a stimulus reconstruction approach to estimate the quality of auditory processing from the EEG. In particular, we focused on the reconstruction of the auditory envelope of the noisy speech from EEG recordings. Our rationale was that participants’ ability to extract relevant cues from noisy speech should be reflected in a better entrainment of EEG activity by the noisy speech (cf. Baltzell et al. 2017) and/or a better encoding of acoustic features, both resulting in a better ability to reconstruct the envelope of the auditory input from EEG recordings. A similar approach was successfully applied to decode attention when participants are exposed to clear speech in a multitalker environment (O’Sullivan et al., 2015; Legendre et al., 2019), or to reconstruct the envelope of noise-masked speech segments (Cervantes Constantino and Simon 2018).

We first extracted the acoustic envelope of the training and test stimuli in the 2-8 Hz band. This band was chosen for its correspondence with syllabic rhythms and the robust entrainment of EEG oscillations with the speech envelope observed in this frequency band (Giraud and Poeppel 2012; Peelle and Davis 2012; O’Sullivan et al. 2015). To do so, we ran the 10.5 s degraded speech as well as the training stimulus through a peripheral auditory model using the standard Spectro-Temporal Excitation Pattern approach (STEP; Leaver and Rauschecker, 2010). The stimuli were first resampled at 22.05 kHz and passed through a bandpass filter simulating outer and middle-ear pre-processing. Cochlear frequency analysis was then simulated by a bank of linear gammatone filters (N=128 filters). Temporal integration was applied on each filter output by applying half-wave rectification and a 100 Hz low-pass 2nd-order Butterworth filter. Next, square-root compression was applied to the smoothed signals and the power in each frequency band was log-transformed. Finally, the auditory envelope was computed by summing the envelope of the 128 gammatone filters and downsampled to 100 Hz.

For each presentation of the (training or test) stimuli, we processed the EEG recordings as follows. ICA-corrected data epochs were re-referenced to the average of all EEG electrodes, bandpass-filtered between 2 and 8 Hz using a two-pass finite impulse response (FIR) filter, and then resampled at 100 Hz. We trimmed the EEG epochs so that the start and end corresponded to the start and end of the stimulus presentation.

We then used the Multivariate Temporal Response Function (mTRF) Toolbox (v2.0; Crosse et al., 2016) for Matlab to build a linear model between auditory and EEG signals from the training session (clear speech). By using an independent part of the experiment compared to test trials, and by using clear speech, we ensured that the model was not affected by our experimental design and represented normal speech processing. EEG data were shifted relative to the auditory envelope from 0 ms to 300 ms in steps of 10 ms (31 time lags), allowing the integration of a broad range of EEG data to reconstruct each stimulus time point. The linear model was optimised to map the EEG signal from each electrode and time lag to the sound envelope. The obtained filter (matrix of weights: sensor ✕ time lags) was then used in the test trials to reconstruct the stimuli.

In the test trials, we used the model trained on clear speech (training set) to reconstruct the envelope of the noisy stimuli. This was done independently for each of the two presentations of the stimuli in each trial. Finally, the reconstructed envelope was compared to the envelope of the degraded stimulus played for this trial (NVS or SWS) by computing the Pearson’s correlation coefficient between the real and reconstructed envelope of the degraded speech. We computed this coefficient for the three repetitions of the same sentence in each stimulus presentation. This coefficient (bounded between -1 and 1) was used as an index of the quality of the stimulus reconstruction. In our analyses, we focused on the first presentation of the sentence within a given trial and the first following the presentation of the correct (P+), incorrect (P-), or no (P0) visual sentence information (first 3.5 s of each presentation). This decision mitigated potential fluctuations in task engagement over the course of the 2nd presentation depending on whether stimuli elicited pop-out.

#### Time-frequency decomposition

EEG data from test trials were subjected to spectral (time-frequency) analysis. Preprocessed datasets were re-referenced to the average of linked mastoids. Spectral power estimates were then computed for epochs spanning -2 to 12 s relative to stimulus onset over a frequency range of 1 to 30 Hz (1 Hz increments) using the ‘ft_freqanalysis’ function (Hanning taper length = 1 s; 100 ms increments). As in the stimulus reconstruction analysis, time-frequency analysis was limited to the first iteration of each sentence presentation period (timepoints spanning [0.5, 3] s; first and last 0.5 s omitted to avoid spurious/confounding effects pertaining to stimulus onset/offset and spectral leakage).

Channel-level spectral power estimates were averaged across time for each trial, and averaged across trials for each factorial combination of sentence type, prior condition, and presentation order. Averaged power estimates were then log_10_ transformed and subjected to a nonparametric cluster-based permutation analysis (Maris and Oostenveld 2007) as implemented in FieldTrip. Briefly, this procedure involved computing dependent-samples *t*-tests across pairwise power estimates for each corresponding channel ✕ frequency bin, identifying *t*-values that exceed a specified alpha threshold (0.025, two-tailed test), and clustering these samples into spatio-spectrally contiguous sets (minimum 2 neighbouring channels located within a 40 mm radius; average 3.9 neighbours per channel). *T*-values within each resolved cluster were then summed and the maximum value assessed against a Monte Carlo simulation-based reference (null) distribution generated over 1000 random permutations. We derived a Monte-Carlo *p*-value from this comparison, which we used to determine the significance of the identified clusters.

To test the interaction of interest, the difference between 1st and 2nd presentation power estimates was contrasted across pairwise combinations of prior conditions for each sentence type. Clusters with a Monte Carlo *p*-value <.05 were deemed indicative of a significant difference between contrasts. Importantly, this procedure only licences inferences about the existence of a statistically significant difference between contrasts; it does not permit the topographic or spectral localisation of such effects (see Maris and Oostenveld, 2007; Maris, 2012; Sassenhagen and Draschkow, 2019). This caveat notwithstanding, the frequency bounds of the resolved clusters were used to inform the selection of frequency band limits in the subsequent linear mixed-effects regression analysis (see *Statistical modelling* section below).

#### Time-resolved oscillatory activity

To examine the temporal evolution of electrophysiological dynamics during sentence processing, we complemented our analysis of time-averaged changes in spectral power with time-resolved profiles of induced (i.e. non-phase-locked) oscillatory activity. These profiles were derived using an intertrial variance method of estimating event-related (de)synchronisation (Kalcher and Pfurtscheller 1995; Pfurtscheller and Lopes da Silva 1999).

ICA-corrected, mastoid-referenced EEG data were high- and low-pass filtered (one-pass zero-phase FIR filters) into the same frequency bands derived following the time-frequency cluster-based permutation analysis. For each frequency band, filtered signals were divided into factorial combinations of stimulus type, prior condition, and presentation order, and the evoked response subtracted from each set. Waveforms were then squared, log_10_ transformed, and averaged within each set. The resulting spectral profiles were smoothed using a moving average filter (‘movmean’, 500 ms sliding window) and downsampled to 10 Hz. Please note, induced responses were not scaled according to a prestimulus reference period, since doing so would have conflated differences between the prior and no-prior conditions during the visual presentation period with differences between these conditions during the 2nd auditory presentation period.

Spectral profiles of induced activity were compared across conditions using a similar cluster-based permutation approach to that described above (*Time-frequency decomposition*), with the exception that channel-wise power estimates were clustered over the temporal (rather than frequency) dimension. Permutation tests evaluating the temporal evolution of induced power dynamics were performed on the entire 2nd presentation window ([-1, 11]s). This analysis was conducted separately for SWS and NVS sentences, in order to explore qualitative differences in the temporal patterning of neural responses to these two stimulus types. The temporal dynamics of pupil size were examined using the same statistical procedure in the temporal domain, but without the spatial (electrode) dimension.

### Statistical modelling

Statistical analysis of trial-level subjective clarity ratings, frequency band power, and stimulus reconstruction scores was performed in *R* (v4.1.1; R Core Team 2021). Our general strategy for each analysis was to fit the appropriate mixed-effects model to the dependent variable of interest from the 2nd presentation, and regress these estimates onto the corresponding estimate from the 1st presentation (including the 1st presentation estimate as a covariate essentially functions as a form of baseline correction; see Alday, 2019). Trial number was also included as a proxy for time-on-task, thereby controlling for the effects of perceptual learning (Davis and Johnsrude 2007; Eisner et al. 2010; Sohoglu and Davis 2016) and other potential sources of nonstationarity (Benwell et al. 2019). Additional independent variables were stimulus type (SWS, NVS), prior condition (P+, P-, P0), and the interaction between these factors, which were introduced into each model in that order. Model comparisons (see below) were performed using the ‘anova’ function to assess whether the additional complexity introduced by each new fixed (and accompanying random) effect term was merited by a sufficient improvement in model fit. Categorical variables (stimulus type, prior condition) were sum-to-zero contrast-coded (reference level coded -1).

All mixed-effects models were fitted with by-participant random intercepts. We attempted to fit maximal random effects structures for all fixed effects of interest (i.e. stimulus type, prior condition, stimulus type ✕ prior condition) on this intercept (Barr et al. 2013). Simpler random effects structures were selected when the maximal model failed to converge, generated a singular fit, or when random effects could be reduced in complexity without significant impairment of model fit (Matuschek et al. 2017). Random intercepts were also specified for sentence items in all models; EEG electrode channel locations were included as random intercepts in the spectral power models only (see Liebherr et al. 2021, for a similar approach).

Subjective clarity ratings following the 2nd presentation were modelled as ordinal data using (logit-linked) cumulative link mixed-effects models (i.e. proportional odds mixed-models). These models were fit via the Laplace approximation using the ‘clmm’ function from the *ordinal* package (v2019.12-10; Christensen 2019) in *R*. No assumptions about the distance between cut-point thresholds were specified.

Linear mixed-effects models for spectral power (averaged over time and frequency bins; first sentence iteration only) and stimulus reconstruction scores (first sentence iteration only) were fitted using the ‘lmer’ function from the *lme4* package (v1.1-27.1; Bates et al., 2015). In addition to the fixed effects described above (which were again introduced in a sequential fashion to enable model comparison), an ordered factor encoding the clarity rating on the 1st presentation was included as a covariate. Model diagnostics were assessed with the aid of the *performance* package (v0.8.0; Lüdecke et al., 2021).

The significance of main effect and interaction terms for each winning model was assessed using likelihood-ratio chi-square tests from Type-II analysis-of-deviance tables obtained via the *RVAideMemoire* package (v0.9-81; Hervé, 2021) for the cumulative link mixed-effects models; equivalent tables were obtained from the *car* package (v3.0-12; Fox and Weisberg, 2019) for the linear mixed-effects models. Significant effects were disambiguated using post-hoc contrasts (Tukey-corrected for multiple pairwise comparisons; Sidak-corrected for pairwise interaction contrasts) obtained from the *emmeans* package (v1.7.1-1; Lenth 2021), which was also used to estimate marginal mean predictions for model visualisation. For completeness, β coefficients (log odds in the case of the cumulative link mixed-models) and standard errors (*SE*s) are reported alongside corresponding analysis-of-deviance results for significant model terms. Model predictions and individual-level estimates were visualised with the aid of the *tidyverse* package (v1.3.1; Wickham et al., 2019).

### Data and code availability

De-identified raw data are openly available on the Open Science Framework: https://osf.io/5qxds. The code used to produce the analyses reported in this manuscript are archived on GitHub: https://github.com/corcorana/SWS_NVS_code

## Results

### Correct prior information evokes perceptual pop-out

We first examined participant- and group-level average clarity ratings to determine if our protocol was successful in eliciting perceptual pop-out (Figure 1C). We found that prior condition had a significant effect on clarity ratings following the 2nd stimulus presentation: including prior condition within the cumulative link mixed-effects model yielded a significant improvement in fit (𝜒^2^(2)=1325, *p*<.001). Allowing prior condition to interact with stimulus type further enhanced model fit (𝜒^2^(2)=7.41, *p*=.025). This model revealed significant main effects of prior condition (𝜒^2^(2)=57.95, *p*<.001; β_P+_=4.21, *SE*=0.27; β_P-_=-2.03, *SE*=0.13) and stimulus type (𝜒^2^(1)=13.71, *p*<.001; β_SWS_=0.18, *SE*=0.69), in addition to the significant interaction of these factors (β_SWS:P+_=-0.29, *SE*=0.10; β_SWS:P-_=0.15, *SE*=0.08).

To interrogate these results, we performed post-hoc contrasts comparing differences in clarity ratings between stimulus types and prior conditions. Clarity ratings were significantly higher following SWS stimuli than NVS stimuli (z-ratio=2.64, *p*=.008). Clarity was significantly higher for P+ than both the P0 (z-ratio=14.81, *p*<.001) and P- (z-ratio=16.10, *p*<.001) conditions, consistent with the experience of perceptual pop-out. Conversely, clarity levels did not significantly differ between P0 and P- (z-ratio=1.00, *p*=.575), confirming that participants needed to be provided with the correct sentence information for pop-out to occur. Interaction contrasts further confirmed that the increase in clarity ratings observed in the P+ condition was stronger for NVS than SWS stimuli (P+ vs. P0: z-ratio=2.51, *p*=.036; P+ vs. P-: z-ratio=2.56, *p*=.032), owing to the lower average clarity of NVS stimuli in the absence of correct prior information (see Fig 1C).

Finally, all participants performed at or near ceiling level when asked to determine if the visually displayed sentence (P+ or P- condition) corresponded to the auditory stimulus (mean performance: 95.8% +/- 1.3 and 95.0% +/- 1.4 for SWS and NVS, respectively). This indicates that participants were almost always able to distinguish whether correct information had been supplied, even when the perceived clarity of the degraded sentence remained low.

### Correct prior information enhances stimulus reconstruction

Having established the efficacy of our prior condition manipulation, we next explored the neurophysiological substrates of the pop-out effect by examining participant- and group-level average reconstruction coefficients (Figure 2A). The mixed-effects model for reconstruction scores revealed a significant main effect of stimulus type (𝜒^2^(1)=23.49, *p*<.001; β_SWS_=0.036, *SE*=0.007), indicating that reconstruction scores, just as clarity ratings, were higher following SWS than NVS stimuli. Importantly, a significant main effect was also observed for prior condition (𝜒^2^(2)=15.97, *p*<.001; β_P+_=0.022, *SE*=0.006; β_P-_=-0.009, *SE*=0.006). In fact, model comparisons indicated that models not including the prior condition effect fitted the data significantly worse (𝜒^2^(2)=15.92, *p*<.001), but including interaction terms (two-way interaction between prior condition and stimulus type; three-way interaction between prior condition, stimulus type, and baseline reconstruction score) did not fit the data significantly better (all *p*s>.10).

**Figure 2:**
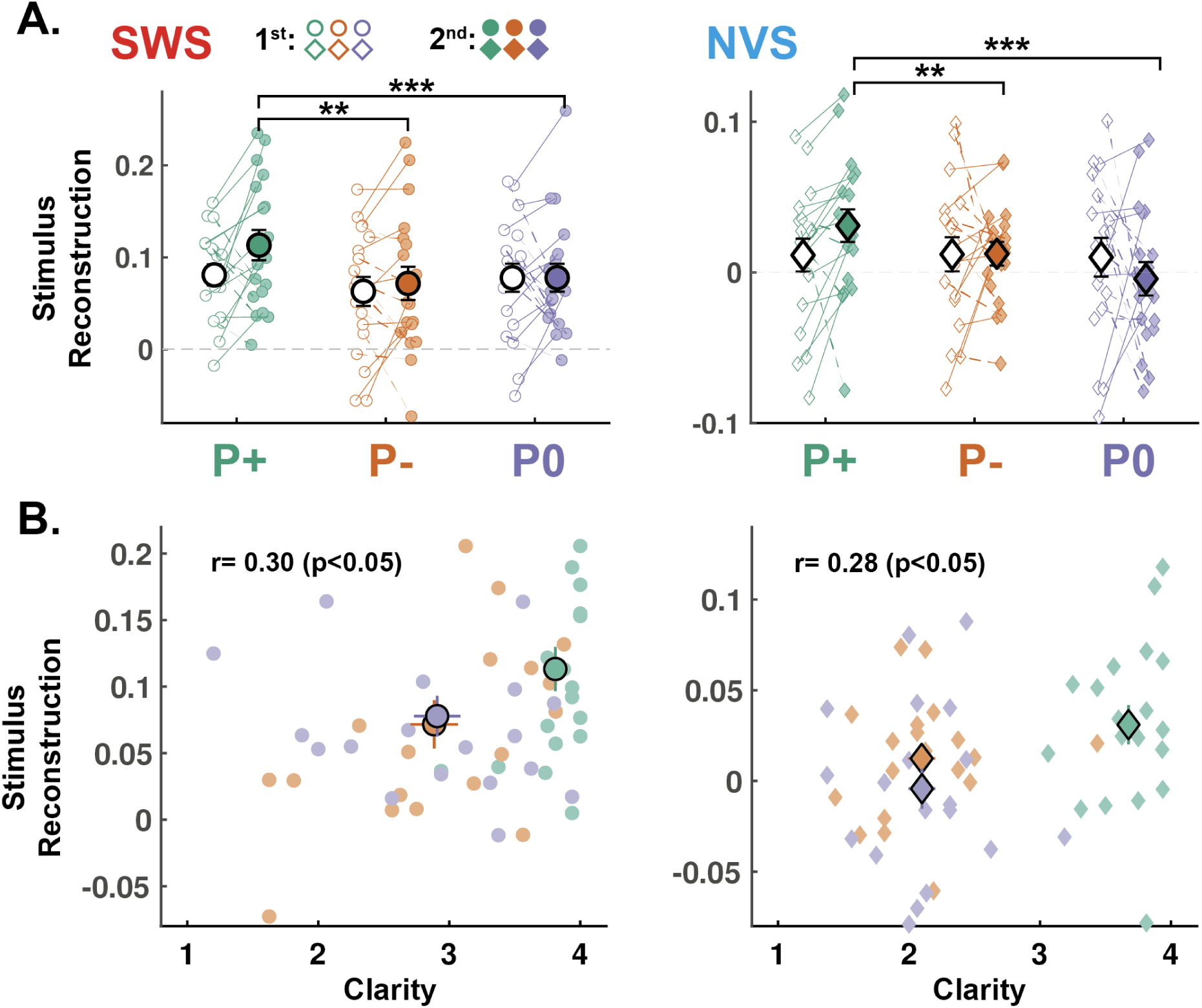
Correct priors improve stimulus reconstruction. A: The envelope of noisy speech was reconstructed from EEG recordings (N=19 participants, see Methods) and a stimulus reconstruction score was computed for the first 3.5 s (1st iteration of the sentence) of each stimulus presentation (1st: unfilled markers; 2nd: filled markers) and for the SWS (left, circles) and NVS (right, diamonds) stimuli separately. Reconstruction scores are averaged for each stimulus type and prior condition (P+: green, P-: orange, and P0: purple). Individual data-points are shown with small circles (SWS) and diamonds (NVS). The two average ratings of each participant and each category are connected with a continuous line if it increases from the 1st to 2nd presentation and a dashed line if it decreases. Large circles and diamonds show the average across the sample (N=19 participants) and error bars show the standard error of the mean (SEM) across participants. Stars indicate the significance levels of post-hoc contrasts across condition levels (***: p<.001, **: p<.01, *: p<.05). B: Correlation between clarity ratings and reconstruction scores on the 2nd presentation for SWS (left, circles) and NVS (right, diamonds). Individual data-points are shown with small circles (SWS) and diamonds (NVS). Large circles and diamonds show the average across the sample (N=19 participants) and error bars show the standard error of the mean (SEM) across participants. The Pearson’s correlation coefficient computed across conditions for the SWS and NVS is shown on each graph along with the associated p-value.

Again, we interrogated the main effect of prior condition with post-hoc pairwise comparisons. These contrasts revealed that reconstruction scores for the 2nd presentation were higher in the P+ compared to P0 (t-ratio=3.66, *p*<.001) and P- (t-ratio=3.22, *p*=.004) conditions, respectively. Reconstruction scores did not significantly differ between P- and P0 (t-ratio=0.44, *p*=0.90). In sum, the effect of prior condition on reconstruction scores matched the pattern of effects found on clarity ratings, suggesting a link between perceptual pop-out and auditory cortical encoding.

We subsequently examined whether the stimulus reconstruction scores could predict the clarity of stimuli on the 2nd presentation above and beyond the prior condition. To do so, we re-fitted the cumulative link mixed-effects model reported above with additional terms encoding the main-effect of reconstruction score, and its interaction with stimulus type and prior condition. These additional terms delivered a significant improvement in model fit (𝜒^2^(6)=13.27, *p*=.039). This model revealed a significant three-way interaction (𝜒^2^(2)=8.43, *p*=.015; β_SWS:P+:REC_=1.51, *SE*=0.59; β_SWS:P-:REC_=-0.24, *SE*=0.47), whereby higher reconstruction scores predicted greater improvement in the subjective clarity of SWS stimuli following the provision of correct vs. no sentence information (z-ratio=2.87, *p*=.012). No such differences were found for the P+ vs. P- (z-ration=1.82, *p*=.191) nor P- vs. P0 contrasts (z-ratio=1.40, *p*=.411).

### Prior information exerts frequency-specific effects on speech processing

We next examined how the provision of correct or incorrect prior information impacted brain responses to degraded speech. Grand-average time-frequency representations from the 2nd presentation period are displayed for each prior condition in Figure 3A. Cluster-based permutation tests revealed significant differences in the average power across prior conditions for each stimulus type. Relative to the absence of written sentence information (P0), receiving correct sentence information (P+) resulted in significant positive clusters (indicative of increased mean power) spanning 12 to 17 Hz in the SWS condition (*p*=.006), and 11 to 15 Hz in the NVS condition (*p*=.005). Similarly, receipt of incorrect sentence information (P-) resulted in a significant positive cluster spanning 10 to 15 Hz in the SWS condition (*p*=.002). No significant clusters were identified for the corresponding NVS contrast. Topographies visualising the distribution of these clusters are presented in Figure 3B.

**Figure 3:**
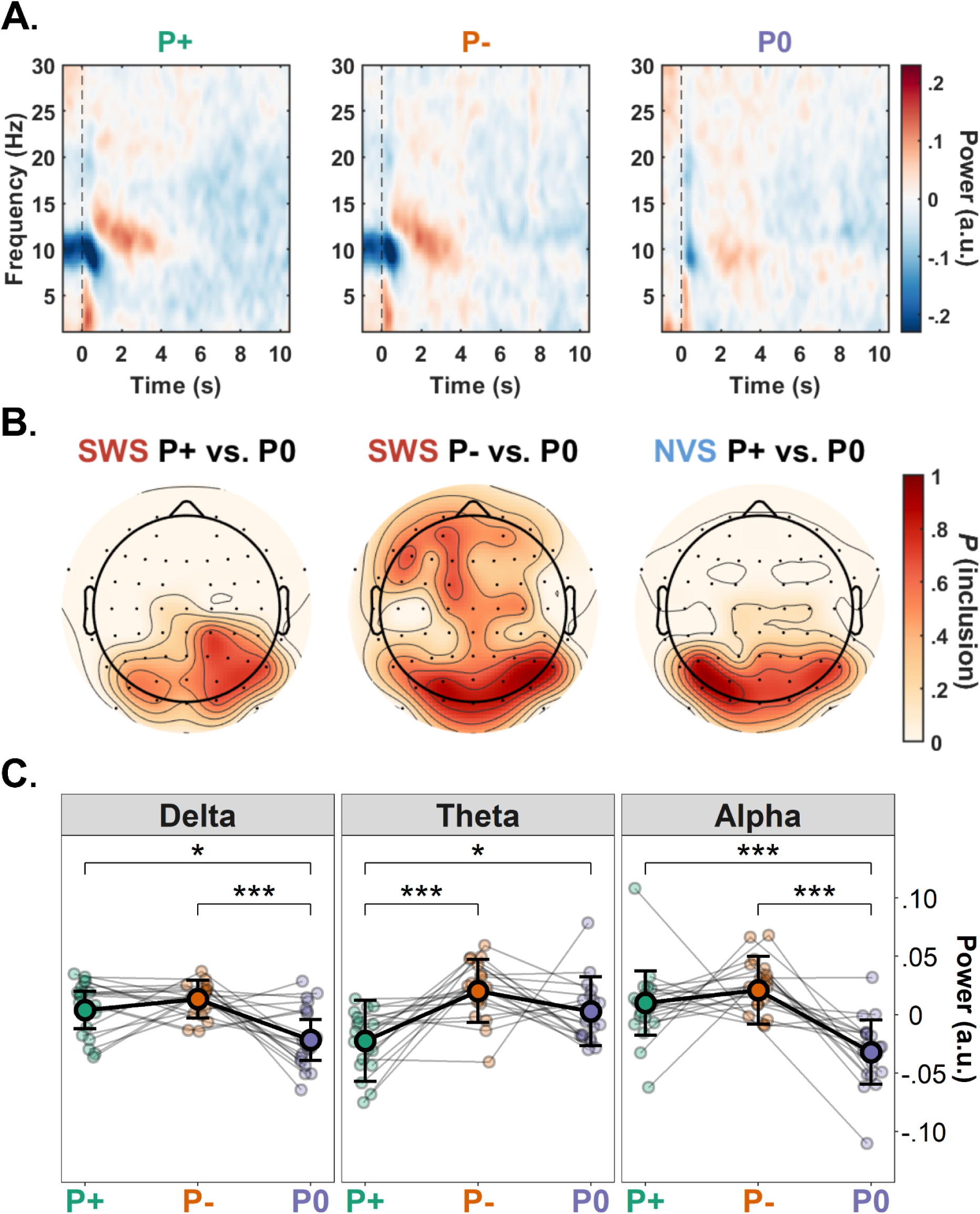
Time frequency analysis and mixed-effects modelling. A: Time-frequency representation depicting grand-average power over the course of the 2nd sentence presentation following the provision of correct (P+), incorrect (P-), or no (P0) written sentence information (averaged across stimulus types). Each presentation comprised 3 iterations of the same noisy stimulus (∼3.5 s each). Spectral power estimates from each frequency bin were baseline-corrected using the mean power estimate from the corresponding frequency bin averaged over all time bins spanning the 1st presentation period. B: Topographic distribution of electrodes’ involvement in the clusters identified via cluster-based permutation analysis of the first sentence iteration. Scale indicates probability of electrode inclusion (i.e. the proportion of times an electrode was included within the cluster) within the 10-15 Hz range used to define the alpha-band. These plots indicate that significant clusters were predominantly composed of electrodes over posterior scalp regions for P+ vs. P0 contrasts, and more broadly distributed for the SWS P- vs. P0 contrast. C: Visualisation of linear mixed-effects model predictions for delta (1-3 Hz), theta (4-9 Hz), and alpha (10-15 Hz) power during the first sentence iteration for each prior condition (P+: green, P-: orange, and P0: purple; averaged across stimulus types). Individual data-points are shown with small circles. Large circles show the estimated marginal means for the prior condition across the sample (N=19 participants); error bars show the standard error of the mean (SEM) across participants. Stars indicate the significance levels of post-hoc contrasts across condition levels (***: p<.001, **: p<.01, *: p<.05). Note, estimates have been mean-centred for the purposes of visualisation.

In order to investigate these modulations in (high) alpha-band activity in more detail, we used linear-mixed effects regression analysis to model trial-level power fluctuations in the frequency band spanning 10 to 15 Hz. Additionally, frequency bands were also specified for the delta (1-3 Hz), theta (4-9 Hz) and beta (16-30 Hz) frequencies. It is worth noting that all these bands have been associated with speech processing in previous studies (see Discussion). All electrode channels were included in the random effects structure for each model. In each set of nested model comparisons across the four frequency bands, the full model (i.e. including the fixed effect and by-participant random slope interactions between stimulus type and prior condition) demonstrated significantly better fits than the reduced models.

The main effect of stimulus type was significant in the delta (𝜒^2^(1)=5.87, *p*=.015; β_SWS_=-0.007, *SE*=0.003) and theta (𝜒^2^(1)=5.53, *p*=.019; β_SWS_=-0.005, *SE*=0.003) models, indicating that NVS stimuli tended to elicit higher mean power than SWS stimuli. This effect was non-significant in the alpha (𝜒^2^(1)=1.78, *p*=.182) and beta (𝜒^2^(1)=1.68, *p*=.194) models. The main effect of prior condition was significant for all frequency bands except beta (Delta: 𝜒^2^(2)=16.63, *p*<.001; β_P+_=0.005, *SE*=0.005; β_P-_=0.015, *SE*=0.004; Theta: 𝜒^2^(2)=15.56, *p*<.001; β_P+_=-0.023, *SE*=0.006; β_P-_=0.020, *SE*=0.005; Alpha: 𝜒^2^(2)=30.70, *p*<.001; β_P+_=0.010, *SE*=0.007; β_P-_=0.021, *SE*=0.006; Beta: 𝜒^2^(2)=0.14, *p*=.931). The interaction between stimulus type and prior condition was significant in the alpha model (𝜒^2^(2)=6.21, *p*=.045; β_SWS:P+_=-0.001, *SE*=0.004; β_SWS:P-_=0.008, *SE*=0.004), but non-significant in the remaining frequency bands (Delta: 𝜒^2^(2)=0.97, *p*=.617; Theta: 𝜒^2^(2)=2.88, *p*=.237; Beta: 𝜒^2^(2)=0.26, *p*=.877).

The estimated effects of prior condition on mean spectral power in the delta-, theta-, and alpha-bands are visualised in Figure 3C. Post-hoc comparisons revealed that delta power was significantly increased following P+ compared to P0 (z-ratio=2.37, *p*=.047), and P- compared to P0 (z-ratio=3.91, *p*<.001); the difference between P+ and P- was non-significant (z-ratio=1.48, *p*=.300). Alpha-band power showed a similar pattern, where power was increased following P+ compared to P0 (z-ratio=3.60, *p*<.001), and following P- compared to P0 (z-ratio=5.47, *p*<.001). Notably, the difference between P- and P0 conditions was more pronounced for SWS than NVS stimuli (z-ratio=2.49, *p*=.038). Again, there was no significant difference in alpha power between P+ and P- (z-ratio=0.95, *p*=.607). By contrast, the theta model revealed a significant decrease in power following P+ compared to P0 (z-ratio=2.64, *p*=.022), and P- (z-ratio=4.27, *p*<.001); the difference between P- and P0 was non-significant (z-ratio=1.94, *p*=.127).

### Prior information induces increased alpha power and pupil size during sentence processing

Finally, we focused more specifically on the effect of prior information (P+ or P- vs. P0) on participants’ neurophysiological activity, regardless of the correctness of this information. Consistent with our analysis of time-averaged spectral power, the time-course of induced alpha-band activity during the 2nd presentation period was modulated by the provision of prior information. Cluster-based permutation analysis across all electrodes confirmed that P+ and P- both induced significant increases in alpha power compared to P0 during the 2nd auditory presentation (Figure 4A). These positive clusters were broadly distributed over the entire scalp, with posterior electrode sites revealing effects that spanned the largest number of time bins (SWS: P+ vs. P0 = [0.8, 3.6]s, P- vs. P0 = [0.8, 3.3]s; NVS: P+ vs. P0 = [1.1, 3.7]s, P- vs. P0 = [0.9, 3.7]s). Note that these effects were preceded by significant negative clusters spanning the pre-stimulus period, most likely reflecting alpha desynchronisation in response to the processing of visually-presented sentence information (SWS: P+ vs. P0 = [-1, 0.4]s, P- vs. P0 = [-1, 0.3]s; NVS: P+ vs. P0 = [-1, 0.6]s, P- vs. P0 = n.s.). Induced activity did not significantly differ between P+ and P- conditions.

**Figure 4:**
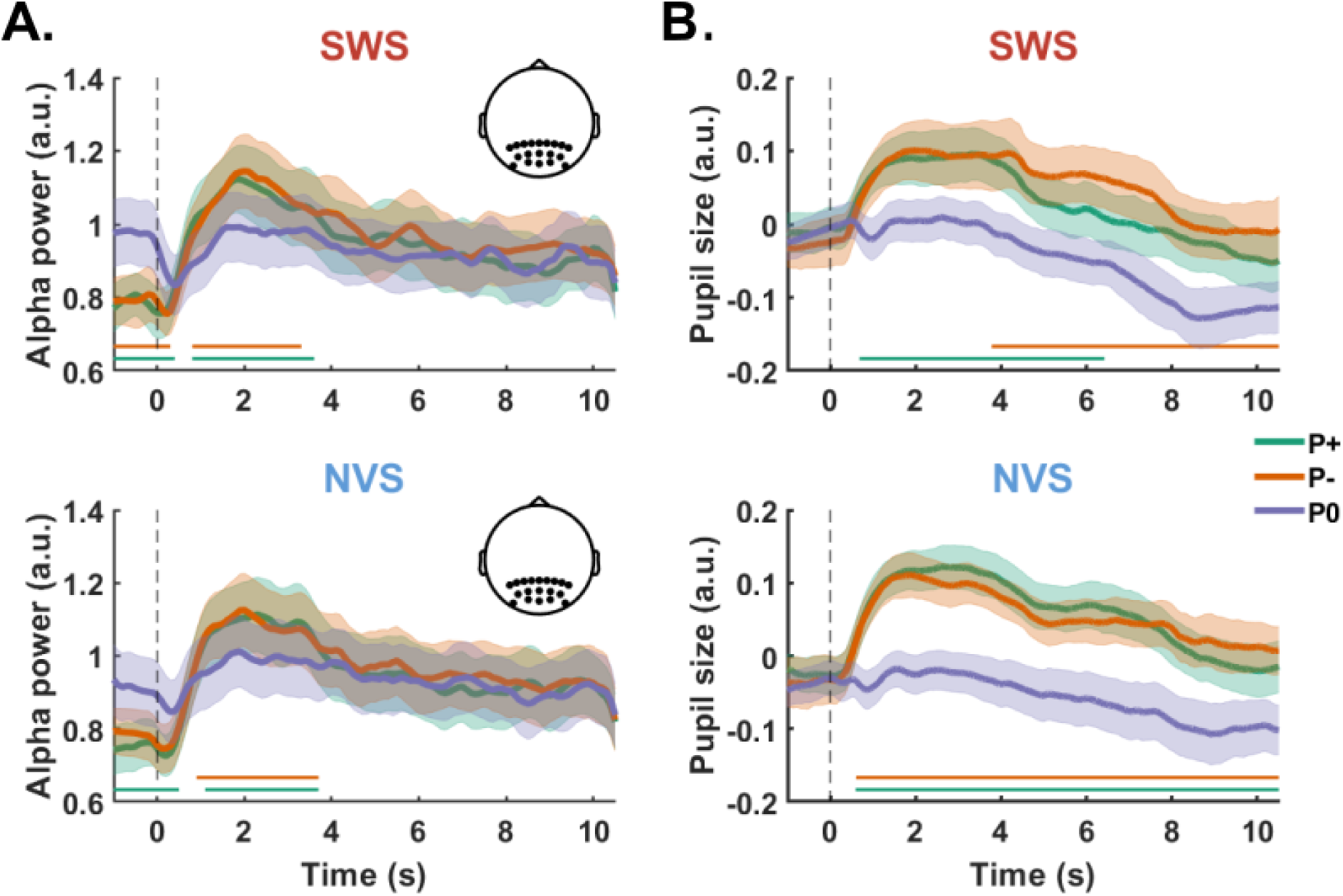
Expectations modulate induced alpha power and pupil size. Temporal dynamics of induced alpha power (A) and pupil size (B) over the course of the 2nd presentation period for each stimulus type (top: SWS; bottom: NVS) and prior condition (P+: green, P-: orange, and P0: purple). Alpha power is averaged over parieto-occipital electrodes (see black dots on the inset) and expressed as log_10_ units. Pupil size is averaged across the two eyes and expressed in arbitrary units. Error shades show the standard error of the mean (SEM) across participants (N=19 participants for alpha power and N=17 for pupil, see Methods). Horizontal bars show the clusters of timepoints showing significant differences (cluster-permutation, p<.05, see Methods) between the P+ and P0 conditions (green), and P- and P0 conditions (orange).

We complemented our analysis of alpha power dynamics with a cluster-based permutation on pupil dilation responses during sentence processing. Similar to the alpha-band findings above, P+ and P- conditions both evoked increased pupil size compared to P0 (Figure 4B). When examining SWS and NVS stimuli separately, we observed a significant cluster for both P+ vs P0 and P- vs P0 contrasts (SWS: [0.7, 6.4]s, [3.8, 11]s; NVS: [0.6, 11]s, [0.6, 11]s, for P+ vs P0 and P- vs P0 respectively).

## Discussion

This study leveraged the pop-out phenomenon to investigate the predictive mechanisms underpinning continuous speech processing. Our key finding suggests that sentence pop-out is mediated via a top-down mechanism that enhances the quality of auditory cortical representations. This observation – which was replicated across two markedly different kinds of acoustic degradation – is consistent with recent electrophysiological evidence that the encoding of degraded speech features is significantly improved after exposure to the undistorted speech stream (Holdgraf et al. 2016; Baltzell et al. 2017; Di Liberto et al. 2018). Crucially, our use of cross-modal (written) information to induce expectations about sentence content ensured these effects could not have arisen due to prior auditory experience of the clear utterance, but were exclusively driven by top-down information. Our design further ensured that the quality and quantity of acoustic stimulation were held constant across conditions, thereby eliminating potential confounds stemming from stimulus novelty and repetition effects. Our data thus provide compelling evidence in support of a predictive coding explanation of sentence pop-out.

This study is the first to use a neural decoding approach to explore the effects of prior information on sentence pop-out. Combining this approach with complementary analyses of spectral power and pupil diameter revealed distinctive patterns of neurophysiological activity that differentiated the content-specific processes (decrease in theta power) associated with the experience of sentence pop-out (P+ condition) from context-specific processes (increase in delta power, alpha power, and pupil size) associated with the generic processing of degraded speech in the context of prior information (P+ and P- conditions). Similar results were observed regardless of whether the acoustic stimulus had been degraded via the removal of spectral or temporal features, speaking to the generality of these effects as indices of auditory perceptual inference. We interpret these findings in light of previous studies of auditory filling-in, speech processing under adverse listening conditions, and the spectral architecture of predictive coding (Arnal and Giraud 2012; Bastos et al. 2012, 2020; Fontolan et al. 2014; Sedley et al. 2016; Auksztulewicz et al. 2017).

### Sentence pop-out is accompanied by enhanced stimulus reconstruction

In line with previous studies using written text to elicit the pop-out of degraded words (e.g., Sohoglu et al., 2012; Sohoglu and Davis, 2016), sentence intelligibility was markedly improved by the provision of correct prior information (P+) only. Incongruent prior information (P-) did not significantly affect clarity ratings compared to the neutral prior (P0) condition (cf. Sohoglu et al., 2014). Although noise-vocoded speech (NVS) was rated less-clear on average than sine-wave speech (SWS), sentence pop-out was reliably obtained across both stimulus conditions.

This pop-out effect was accompanied by two main electrophysiological correlates: (1) improved stimulus reconstruction (an index of cortical speech envelope tracking), and (2) theta-band (4-9 Hz) power suppression. The stimulus reconstruction finding suggests that information extracted from the written sentence promotes the modulation of low-frequency EEG activity while listening to the corresponding sentence, such that the phase dynamics of the EEG signal better approximate the temporal fluctuations of the speech envelope. This finding is striking for at least two reasons: First, the participant never hears the undistorted version of the sentence at any point in the experiment; hence, the effect is likely mediated by top-down mechanisms rather than low-level adaptations induced by prior sensory experience. Second, the decoding model used to reconstruct the original speech envelope was trained on brain responses to natural, continuous speech (i.e. the audiobook). As such, the model was never exposed to the particular acoustic features of degraded stimuli, suggesting that improvements in stimulus reconstruction could be detected on the basis of generalisation from cortical responses to clear speech.

The enhancement of stimulus reconstruction quality in the P+ condition, and the additional improvement in goodness-of-fit brought about by the incorporation of trial-level reconstruction scores into the clarity rating model, support the notion that reconstruction quality is a reliable indicator of subjective speech clarity. Given that clarity and reconstruction scores were both improved by correct sentence information in the absence of any physical alteration of the auditory stimulus, our findings suggest that top-down information contributes to the experience of pop-out by restoring or enhancing the spectro-temporal detail of degraded speech features (cf. Holdgraf et al. 2016; Cervantes Constantino and Simon 2018). This observation accords with previous reports that the quality of speech tracking covaries with speech intelligibility (Ahissar et al. 2001; Luo and Poeppel 2007; Gross et al. 2013; Peelle et al. 2013; Ding and Simon 2014; Doelling et al. 2014), as when vocoded speech is rendered intelligible after hearing the clear version of the utterance (Baltzell et al. 2017). These results also complement findings from studies that directly manipulated speech tracking and intelligibility through top-down attentional modulation (Rimmele et al. 2015) and bottom-up transcranial stimulation (Riecke et al. 2018).

### Theta-band suppression indexes sentence comprehension

Improved sentence comprehension and stimulus reconstruction in the P+ condition were accompanied by a relative reduction in theta-band activity. Mean theta power was also more reduced while listening to SWS stimuli, which tended to elicit higher clarity ratings on average than NVS stimuli (suggesting that NVS constituted a more acoustically (or cognitively) challenging stimulus than SWS; see Peelle, 2018). These findings are consistent with previous reports linking perceptual filling-in and speech intelligibility with the attenuation of sensory cortical responses (e.g., Sohoglu et al., 2012; Sohoglu and Davis, 2016), including evidence directly implicating theta-band suppression (e.g., Riecke et al., 2009, 2012; Strauß et al., 2014a). They also highlight the utility of complementing stimulus reconstruction with spectral power analysis.

The inverse relation between theta-band power and speech intelligibility can be explained by the involvement of theta dynamics in the retrieval and integration of linguistic representations during online sentence processing (Bastiaansen et al. 2010; Halgren et al. 2015; Lam et al. 2016; Piai et al. 2016; Cross et al. 2018). From a predictive coding perspective, the provision of correct prior information furnishes the listener with an accurate prediction (hypothesis) about the auditory input they are about to encounter. Such information activates lexical representations in working memory, engendering top-down messages that propagate through descending neuronal pathways to sensory cortices. In this way, correct priors may instantiate neural ‘templates’ that facilitate the extraction and integration of syntactic and phonological structures from the acoustic stream (Hickok et al. 2011; Tuennerhoff and Noppeney 2016), thereby inhibiting theta-band activity (cf. Keitel et al., 2017; Rommers et al., 2017; Donhauser and Baillet, 2020).

### Delta power, alpha power, and pupil dilation as correlates of active listening

Listening to degraded speech following the provision of written sentence information (P+, P-) resulted in elevated delta-band power compared to the absence of such information (P0; Figure 3C). Delta oscillations have been implicated in the synthesis of higher-level linguistic structure (Keitel et al. 2018; Meyer and Gumbert 2018; Molinaro and Lizarazu 2018; Etard and Reichenbach 2019; Kaufeld et al. 2020), although much of this literature concerns phase (rather than power) dynamics. Elevated delta-band power might derive from attempts to parse continuous speech according to the segmentation prescribed by the written sentence (cf. Ding et al., 2016; Bonhage et al., 2017; Meyer et al., 2017). Alternatively, it might reflect increased phase-synchrony driven by precise expectations about the timing of salient auditory input (Lakatos et al. 2008; Schroeder and Lakatos 2009; Calderone et al. 2014; Arnal et al. 2015; Kayser et al. 2015; Boucher et al. 2019).

Increased alpha-band power and pupil dilation have received comparatively more widespread attention in the speech comprehension literature, most notably in association with effortful, ‘active’ listening under adverse conditions (Zekveld et al. 2010; Wöstmann et al. 2015; Dimitrijevic et al. 2017, 2019). Parametric increases in alpha-band power (e.g., Obleser and Weisz, 2012; cf. Hauswald et al., 2020) and pupil size (e.g., Winn et al., 2015; cf. Zekveld et al., 2018) have been reported as the severity of speech degradation intensifies, complementing recent evidence that covariation between pupil diameter and alpha-band activity indexes fluctuations in arousal and attentional states (Pfeffer et al. 2021; cf. Podvalny et al. 2021; Sharon et al. 2021). However, the differences we observed in these dependent variables between the prior and no-prior conditions cannot be ascribed to stimulus properties, since the degree of acoustic degradation was held constant across conditions. Likewise, such differences cannot be explained by sentence (un)intelligibility (cf. Becker et al., 2013) or prior (in)congruence, given the similarity of the responses induced by P+ and P- conditions.

Following previous work implicating alpha oscillations in the top-down inhibition of task-irrelevant cortical networks (Klimesch et al. 2007; Jensen and Mazaheri 2010) or stimuli (Kerlin et al. 2010; Strauß, Wöstmann, et al. 2014; Wöstmann et al. 2016; Wöstmann, Lim, et al. 2017), the transition from alpha desynchronisation during the presentation of written sentence information, to a marked and widespread pattern of synchronisation following auditory stimulus onset, might reflect the dynamic re-allocation of attentional resources from the visual to the auditory domain (i.e. a covert attentional switch from visual sampling during reading to auditory sampling during active listening; cf. Foxe et al., 1998; Fu et al., 2001; Henry et al., 2017). Note that the event-related synchronisation observed in the P+ and P- conditions significantly exceeded corresponding power estimates in the P0 condition (Figure 4A), implying that speech onset promoted a concerted synchronisation of alpha-band oscillations (rather than a mere restoration of baseline levels of activity following the offset of visual stimuli).

An additional (but not mutually exclusive) explanation of this effect derives from the putative role of alpha-band synchronisation in the working memory processes responsible for mapping online auditory inputs to specific linguistic predictions (i.e. hypotheses about sentence content induced by prior information; cf. Obleser et al., 2012; Meyer et al., 2013; Sedley et al., 2016; Wilsch and Obleser, 2016). This explanation is appealing given the close correspondence between the time-course of the alpha-band synchronisation and the first sentence iteration. Given the immediacy of the pop-out effect, attempts to match the prior with incoming acoustic information are unlikely to persist beyond the first sentence iteration -- either the hypothesis instantiated by the prior is confirmed and the sentence correctly parsed (cf. Friston et al., 2021), or it is disconfirmed and abandoned. This hypothetical process would seem concordant with the temporal evolution of the induced response, which peaks ∼2 s following sentence onset, declining thereafter.

While pupil diameter also showed a marked increase at the beginning of the auditory stimulus presentation following written sentence information, this response decayed at a much slower rate than that of the alpha synchronisation. Moreover, the profile of the pupil response following prior information varied across stimulus types: while pupil dilation was protracted in both the P+ and P- conditions during NVS, this effect was curtailed for P+ and delayed for P- during SWS. Such differences cohere with the notion that pupil diameter and alpha-band dynamics may tap dissociable cognitive processes (cf. McMahon et al. 2016; Miles et al. 2017; Alhanbali et al. 2019; Podvalny et al. 2021).

Given the tight linkage between pupil size and the neuromodulatory regulation of arousal and attention (Joshi and Gold 2020; Dahl et al. 2022), we interpret these results in terms of generic aspects of cognitive task engagement (Hess and Polt 1964; Kahneman and Beatty 1966; Beatty 1982; Zekveld and Kramer 2014; Hjortkjær et al. 2020). Increased pupil diameter in both the P+ and P- conditions is consistent with a greater allocation of cognitive resources to the auditory stream when prior information is available, as compared to an absence of prior information in the P0 condition. Participants may have been more quick to disengage from SWS stimuli during the P+ condition since it may have been easier to arbitrate the congruence of the written and acoustic stimuli given the increased clarity of SWS during the 1st auditory presentation. NVS, on the other hand, may have presented a more challenging stimulus in general (cf. lower average clarity ratings; Figure 1C), thus explaining the more consistent and protracted dilation response observed across P+ and P- conditions.

An important caveat to the interpretation of these results is that the low-level visual properties of written text stimuli were not precisely matched to ensure parity across conditions. It is thus feasible that differences in luminance might be partially responsible for the differential patterns of alpha-band power and pupil size dynamics observed during the 2nd presentation period. However, while posterior alpha-band activity may ‘rebound’ following the offset of visual stimuli, such effects typically unfold over the course of ∼1 s (see, e.g., Pfurtscheller and Lopes da Silva 1999). By contrast, the synchronisation effect observed here persisted for 3-4 s, consistent with the duration of the first sentence iteration. Moreover, the evoked pupil response in the period preceding visual stimulus offset was very similar across conditions, implying that the average difference in luminance between P0 and P+/P- conditions was negligible (recall that the pupil response was baseline-corrected to the pre-trial interval, which always featured a blank screen). Finally, we note that the interaction between stimulus type and prior condition in the alpha power model, and the distinct patterning of pupil responses >4 s after the onset of SWS stimuli, are difficult to explain in terms of visual stimulus differences. Rather, these specific response profiles are more likely to derive from the cognitive factors outlined above.

## Conclusion

This study isolated the electrophysiological effects of cross-modal prior information on auditory cortical sentence processing, providing further evidence of the top-down predictive mechanisms that support continuous speech comprehension. By manipulating the content of prior expectations while holding bottom-up auditory input and prior stimulus exposure constant, we found that correct expectations led to improved perceptual clarity and enhanced stimulus reconstruction, which could result from the enhanced cortical representation of degraded speech. Furthermore, neurophysiological measures revealed that these effects were accompanied by distinctive functional profiles: while theta activity was relatively reduced following correct sentence information only, delta power, alpha power, and pupil size were all increased following *any* written information. These findings suggest that theta-band activity indexes the efficiency of incremental sentence processing and is sensitive to speech intelligibility, whereas delta- and alpha-band oscillations, along with pupil size, may track more domain-general predictive mechanisms involved in active listening.

## Acknowledgments

We thank Madeline Daveney and Jing Xuan Wang for their help with data collection. We also thank four anonymous reviewers for feedback that significantly improved the quality of the manuscript.

